# Identification of novel PGRP3 protein -protein interactions using yeast two hybrid system

**DOI:** 10.1101/2022.12.16.520769

**Authors:** Li-jung Lin, Clarence Lee, Srivathsan Ranganathan, Jens Kreth, Justin Merritt, Sivaraman Prakasam

**Affiliations:** Department of Periodontology. S.O.D., Oregon Health & Science University, 2730 S Moody Ave, Portland, OR-97201; OHSU Knight Cancer institute, S.O.M., Oregon Health & Science University, 2720 S Moody Ave, Portland, OR-97201; Department of Restorative Dentistry, School of Dentistry, Oregon Health & Science University, 2730 S Moody Ave, Portland, OR-97201; Department of Periodontology. School of Dentistry, Oregon Health & Science University, 2730 S Moody Ave, Portland, OR-97201

**Keywords:** Peptidoglycan recognition Proteins, PGRP, Innate immunity, Yeast two hybrid assay, Co-immunoprecipitation

## Abstract

**Background:** Peptidoglycan recognition protein-3 (PGRP3) is a pattern recognition receptor that binds peptidoglycan to elicit an immune response. How PGRP3 mediates these immunomodulatory effects remains unknown. Identifying the proteins that interact with PGRP3 will help begin to address this important knowledge gap. Therefore, in this study our objective was to identify and validate novel PGRP3-protein interactions.

**Methods:** PGRP3-protein interactions were identified using a yeast two-hybrid system. PGRP3 cloned into a pGBKT7 DNA-BD vector and transformed into a Y2HGold yeast strain served as the bait. A normalized universal human cDNA library cloned in a pGADT7 AD vector and pre-transformed into an Y187 yeast strain served as the prey. Y2HGold and Y187 yeast strains were mated and diploids were plated on selective media. True positive clones were sent in for DNA sequencing and corresponding proteins were identified through NCBI Blast. Two proteins were selected for validation. His-tagged PGRP3 expressing 293T cells were then co-transfected with expression vectors containing the identified proteins tagged with c-Myc. Lysates from co-transfected cells were subjected to anti-His and anti-Myc co-immunoprecipitation and analyzed with western blots.

**Results:** Four unique proteins— MBNL3, RBP5, ATXN2, and GPATCH8—were identified from our yeast two-hybrid screen. Co-transfection and co-immunoprecipitation assays successfully confirmed MBNL3 and RBP5 as positive interactors with PGRP3.

**Conclusions:** Based on the known functions of MBNL3, RBP5, ATXN2, and GPATCH8, we hypothesize that PGRP3 may be involved in RNA processing, endocytic trafficking, and PPARγ pathways related to innate immunity.

## Introduction

Peptidoglycan recognition protein 3 (PGRP3) is an intermediate length isoform of the PGRP family of pattern recognition receptors.^1^ PGRP3 is expressed in skin epidermis, hair follicles, sebaceous and sweat glands, ciliary body of the eye, corneal epithelium, submandibular salivary gland, mucus-secreting glands in the throat, tongue and esophagus squamous epithelial cells, acid-secreting parietal cells (PGRP3)and small and large intestine columnar absorptive cells.^2^ PGRP3 levels are upregulated by bacteria in keratinocytes, fibroblasts, and oral epithelial cells through activation of TLR2, TLR4, Nod1, and Nod2^1,3^. PGRP3 is elevated in saliva and oral tissues during periodontitis.^2^

PGRP3 has antibacterial and anti-inflammatory properties. The dextran sodium sulfate-induced colitis murine model has been used to demonstrate the anti-inflammatory effects of PGRP3. PGRP3 knockout mice were observed to have increased intestinal bleeding, higher apoptosis of colonic mucosa, increased expression of cytokines and chemokines, altered gut microbiota, and decreased survival rates.^4^ Another study showed that PGRP3 knockout mice displayed a more inflammatory gut microflora, increased production of interferon-γ, higher expression of interferon-inducible genes, and a larger number of NK cells in the colon.^5^ PGRP3 interferes with IFN-γ activity in the gut and limits TH17 over activation by promoting Treg accumulation in the skin.^5,6^

Human epithelial colorectal adenocarcinoma cells (Caco-2), which produce cytokines in response to bacterial peptidoglycan, have also been used to examine PGRP3. In a study with colonic pathogens (*S. aureus*), opportunistic pathogens (*M. luteus*), and non-pathogens (*B. subtilis* and *L. rhamnosus*), pro-inflammatory cytokines like IL-12, IL-8, and TNF-α were upregulated in response to all peptidoglycan sources. When PGRP3 was knocked out, the expression levels of these peptidoglycan-induced cytokines increased, suggesting the anti-inflammatory effect of PGRP3.^7^ Another study by the same research group determined that the expression of PGRP3 was dependent on Peroxisome proliferator-activated receptor γ (PPARγ) and its lipophilic ligands.^8^ Oligosaccharides (α3-sialyllactose and Raftilose p95), which modulate the intestinal microbiota to be less inflammatory, were shown to induce activation of PPARγ, thus upregulating the production of PGRP3 to reduce intestinal inflammation.^9^

Thus, there is a preponderance of evidence suggesting that the presence of PGRP3 results in dampening of inflammatory responses. However, the precise mechanisms by which PGRP3 modulates inflammation remains unknown. More importantly, the host molecules that PGRP3 interacts with to bring forth its modulating function is not known. A bioinformatics search of interactomes databases revealed that, per Biological General Repository for Interaction Datasets (BioGRID), PGRP3 has putative interactions with two proteins—Dehydrogenase E1 and Transketolase Domain Containing 1 (DHTKD1) and MHC Class I Polypeptide-Related Sequence A (MICA).^10^ These were identified via affinity capture-mass spectroscopy, however these interactions have not been experimentally confirmed.

Identification and confirmation of the proteins that PGRP3 interacts will reveal the mechanisms and signaling pathways that PGRP3 use to modulate inflammation. Here, we used a GAL4 yeast 2-hybrid approach to identify PGRP3 interacting proteins. Using this method we report the identification of four novel PGRP3-protein interactions. We selected two of these proteins to validate our results using co-immunoprecipitation assays.

## Materials and Methods

### Construction of the bait vector

Human PGRP3 cDNA (Sino Biological) encoding a 357 residue peptide was amplified via polymerase chain reaction using the forward primer 5’-C ATG GAG GCC GAA TTC ATG GCC TTC TTC ATT CTG GGT CTC and reverse primer 5’-G CAG GTC GAC GGA TCC TCA GTG CTT GAA ATG AGG CCA GGT. The secretion signal motif at the N-terminus of PGRP3 was intentionally removed during PCR amplification. After transforming into *E. coli*, the amplified PGRP3 cDNA was digested with BamHI and EcoRI and ligated to a pGBKT7 vector containing the GAL4 DNA-binding domain (BD). Successful recombinant pGBKT7-PGRP3 bait construction was verified with restriction digestion (BamHI, EcoRI) and gel electrophoresis.

### Yeast strains

The yeast strains used were *S. cerevisiae* Y187 (Takara Bio) and Y2HGold (Takara Bio). Additional information regarding their genotype and reporters are provided in Table 1.

**Table 1.**
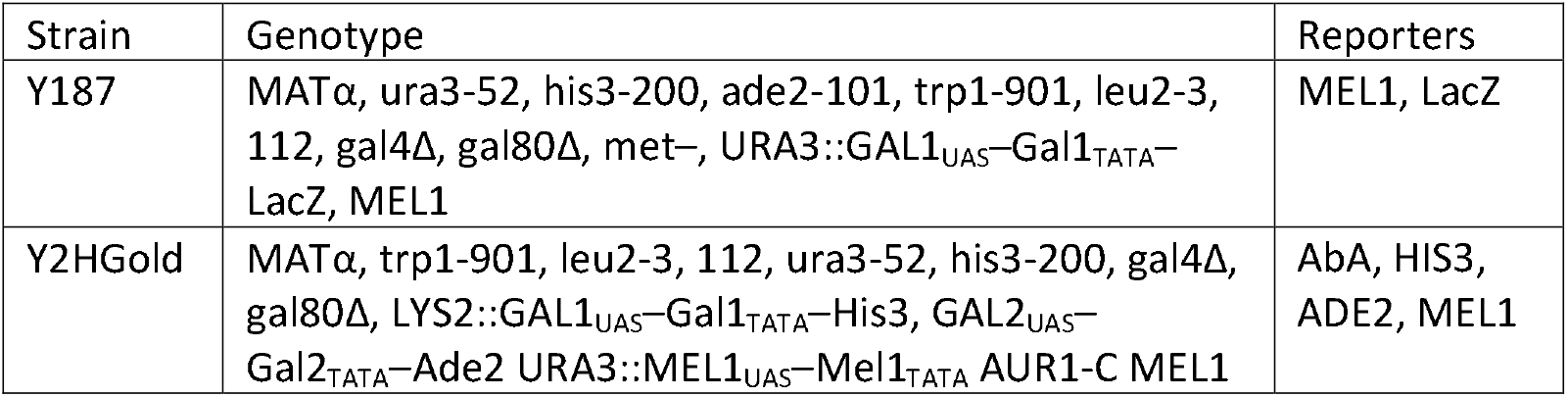
Description of Y187 and Y2HGold genotype and reporters

### Expression of bait protein in yeast

The recombinant pGBKT7-PGRP3 bait construct was transformed into yeast strain Y2HGold via the lithium acetate method per manufacturer protocol (Takara Bio). Transformants were cultured on SD/-Trp (SDO) media for 3-5 days. A representative clone was selected and inoculated into yeast culture medium until the OD_600_ reached 0.5. Proteins were extracted and separated with SDS-PAGE and detected with Western blot using anti-Myc staining.

### Autoactivation and toxicity testing

It was important to ensure that introduction of the pGBKT7-PGRP3 construct did not autoactivate transcription of the reporter gene. Also, it was critical to verify that the bait construct was not toxic to our yeast strains. Transformants were plated on SD/-Trp (SDO), SD/-Trp/X-α-Gal (SDO/X), and SD/-Trp/X-α-Gal/Aureobasidin A (SDO/X/A) for 3-5 days. An empty pGBKT7 vector served as the control. Absence of autoactivation of the bait construct was confirmed through the presence of white colonies on SDO, white colonies on SDO/X, and absence of colonies on SDO/X/A. Absence of toxicity of the bait construct was confirmed by comparing the similar colony size and appearance to that of the control.

### Yeast two-hybrid screening

Yeast two-hybrid screening was performed using a GAL4-based system (Matchmaker Gold Yeast Two-Hybrid System; Takara Bio, Mountain View, CA, USA). Assays and screenings were performed according to the manufacturer’s provided user manual. A Universal Human (Normalized) Mate & Plate^™^ cDNA Library in a pGADT7 vector with the GAL4 DNA-activation domain (AD) and pre-transformed into yeast strain Y187 was acquired from Takara Bio. Mating between Y2HGold transformed with pGBKT7-PGRP3 and Y187 transformed with pGADT7-Universal Human cDNA Library was performed. Presence of diploids were confirmed at 20 hours and subsequently plated on SD/-Ade/-His/-Leu/-Trp supplemented with X-α-Gal and Aureobasidin A. 1:10,000 dilutions of the mated culture were also prepared on SD/-Leu, SD/-Trp, and SD/-Leu/-Trp to calculate mating efficiency. cDNA library plasmids that expressed positive interactions were rescued in *E. coli*. True positive interactions were distinguished from false positive interactions by co-transforming either an empty pGBKT7 vector or a recombinant pGBKT7-PGRP3 vector with a rescued cDNA library pGADT7 vector in Y2HGold competent cells. Cultures were plated on either SD/-Leu/-Trp/X-α-Gal or SD/-Ade/-His/-Leu/-Trp/X-α-Gal/AbA.

### Analysis of positive clones

Confirmed cDNA clones were sequenced using an ABI 3730*xl* 96-capillary DNA Analyzer (Applied Biosystems, Foster City, CA, USA) and a homology search was conducted against standard *Homo sapiens* database sequences using the nucleotide BLAST algorithm on NCBI (National Center for Biotechnology Information).

### Co-immunoprecipitation to confirm protein interactions

293T cells were transfected with a human PGRP3 vector. Stable cell lines expressing His-PGRP3 (293T) were grown on 10 cm plates and transfected with Myc-FLAG-RBP5 or Myc-FLAG-MBNL3 the next day. 48 hours later the cells were harvested. Formaldehyde in-cell crosslinking was performed prior to IP. Cells from one 10 cm plate were suspended in 1 ml PBS containing 1 % formaldehyde and incubated at room temperature for 10 min with gentle agitation. The suspension was spun for 3 min at 1800 g at room temperature and the supernatant was discarded. The pellet was washed with 1 ml of 1.25 M glycine in cold PBS for 5 min at room temperature to quench the crosslinking reaction. The pellet was further washed in PBS, lysed with 250 μl lysis buffer and protease/phosphatase inhibitors at 4°C and sonicated in a water bath sonicator (Fisher) at level 2 for 4 cycles (15 s on/30 s off). The lysates were first cleared by spinning at 16,000g at 4°C for 15 min to remove cell debris, 3 mg of total protein was used for immunoprecipitation with 50μg of anti-His or anti-Myc antibody.

His tag-IP was performed using anti-His affinity resin (GenScript L00439-1) and Myc tag-IP was performed using anti-Myc affinity resin. IPs were performed at 4°C for 2 h and the resin were washed six times in IP buffer. The proteins were eluted with SDS sample buffer and heated at 95°C for 5 min for regular IP samples and heated for 20 min for formaldehyde crosslinking IP samples. The samples were then run on the Bolt 4–20% Bis-Tris Plus Gel (GenScript) and western blot was performed. For western blot analysis, proteins were transferred via a semi-dry procedure on polyvinylidene difluoride membranes (Pall Corporation), blocked for 1 h at RT with 5% milk powder in PBST (PBS, 0.1% Tween 20) and incubated with mouse anti-His or mouse anti-Myc antibody overnight. Membranes were then incubated with goat anti-mouse Alexa Fluor 680 (both 1:10000) for 1 h at RT. Membranes treated with anti-mouse Alexa Fluor 680 were scanned with a fluorescence scanner (Odyssey, LICOR) using excitation/emission wavelengths of 700 and 800□nm.

## Results

### Protein expression of PGRP3 in Y2HGold transformed with pGBKT7-PGRP3 vector

Human PGRP3 cDNA was cloned into a pGBKT7 vector. A restriction enzyme digest assay using BamHI and EcoRI showed two distinct bands at 8.3k bp (pGBKT7) and ~1k bp (PGRP3) (Fig 1a). The resulting pGBKT7-PGRP3 vector was then transformed into yeast strain Y2HGold. Because the pGBKT7 vector includes a c-Myc epitope tag, a Western blot using a c-Myc antibody was performed to confirm PGRP3 expression at ~38kDa (Fig. 1b). An autoactivation test was performed to ensure that the introduction of the pGBKT7-PGRP3 vector did not constitutively activate reporter genes. If the bait vector autoactivated the GAL4 transcription factor, its AUR1-C gene that confers resistance to Aureobasidin A (AbA) would have allowed colonies to grow in plated cultures containing SD/-Trp/X-α-Gal/AbA (SDO/X/A). No colonies were observed, confirming the absence of autoactivation. Also, the bait vector was confirmed as nontoxic to Y2HGold by the presence of colony formation on SD/-Trp media (Fig 1c).

**Figure 1.**
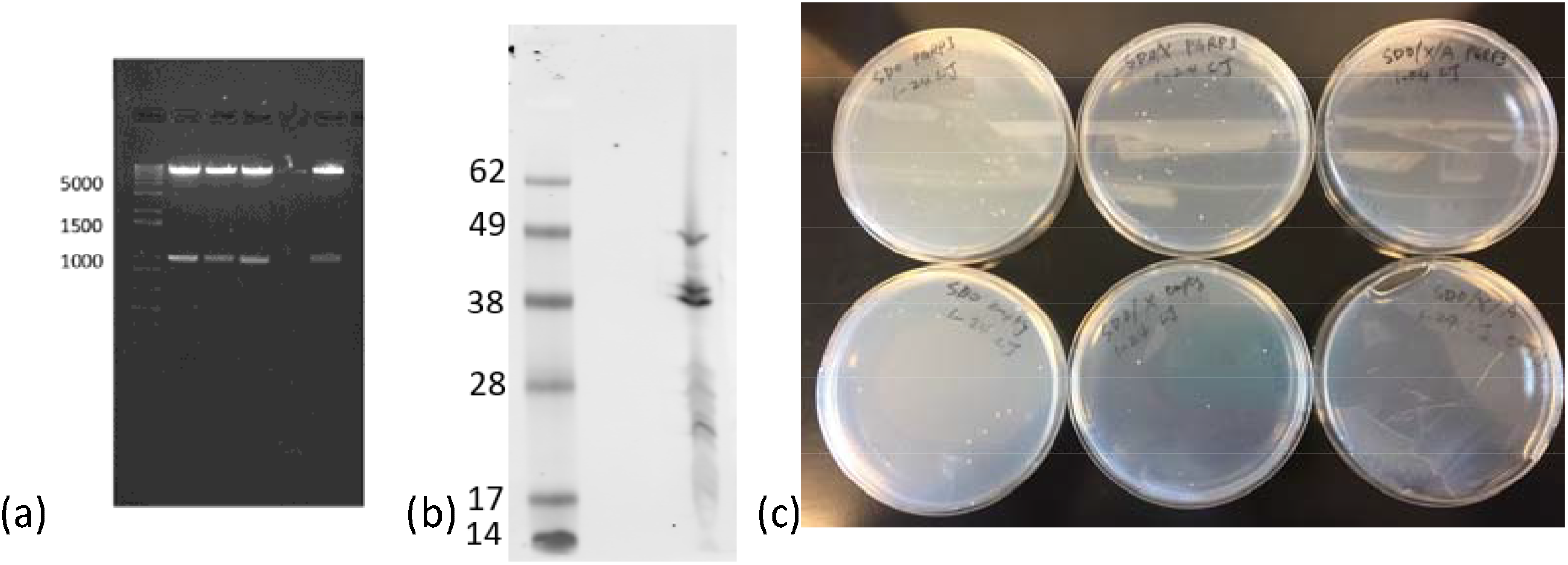
Construction of testing of PGRP3 bait vector (A) Restriction digest with BamHI and EcoRI following introduction of PGRP3 (~1k bp) into pGBKT7 vector (8.3k bp). (B) Western blot with c-Myc antibody after Y2HGold yeast transformation with pGBKT7-PGRP3 construct (Lane 2) and negative PGRP3 control (Lane 1). (C) Y2HGold transformed with pGBKT7-PGRP3 and plated on SD/-Trp (SDO), SD/-Trp/X-α-Gal (SDO/X), and SD/-Trp/X-α-Gal/AbA (SDO/X/A).

### Genes identified from yeast two-hybrid screen

Y2HGold containing pGBKT7-PGRP3 and Y187 containing pGADT7-Universal Human cDNA Library were successfully mated after 20 hours. Plated as a 1/10,000 dilution, approximately 2,000 colonies grew on SD/-Trp, 300 colonies grew on SD/-Leu, and 66 colonies grew on SD/-Trp/-Leu. Between the prey and the bait vectors, the prey was determined to be the limiting factor. Therefore, mating efficiency was calculated to be 22% (viability of diploids ÷ viability of prey library x 100% - 66×10^4^ cfu/mL ÷ 300×10^4^ cfu/mL x 100%).

Diploid colonies were replated on SD/-Trp/-Leu/X-α-Gal/AbA (DDO/X/A). Out of the original 66 colonies, only 4 were viable and could hydrolyze X-α-Gal to yield a blue color (Fig 2). These 4 positive clones were confirmed as genuine by subjecting them to a more stringent media SD/-Trp/-Leu/-Ade/-His/X-α-Gal/AbA (QDO/X/A). Sequencing of positive cDNA inserts identified the corresponding PGRP3-interacting proteins as Muscleblind-like Protein 3 (MBNL3), Retinol Binding Protein 5 (RBP5), Ataxin 2 (ATXN2), and G Patch Domain-containing Protein 8 (GPATCH8) (Table 1).

**Figure 2.**
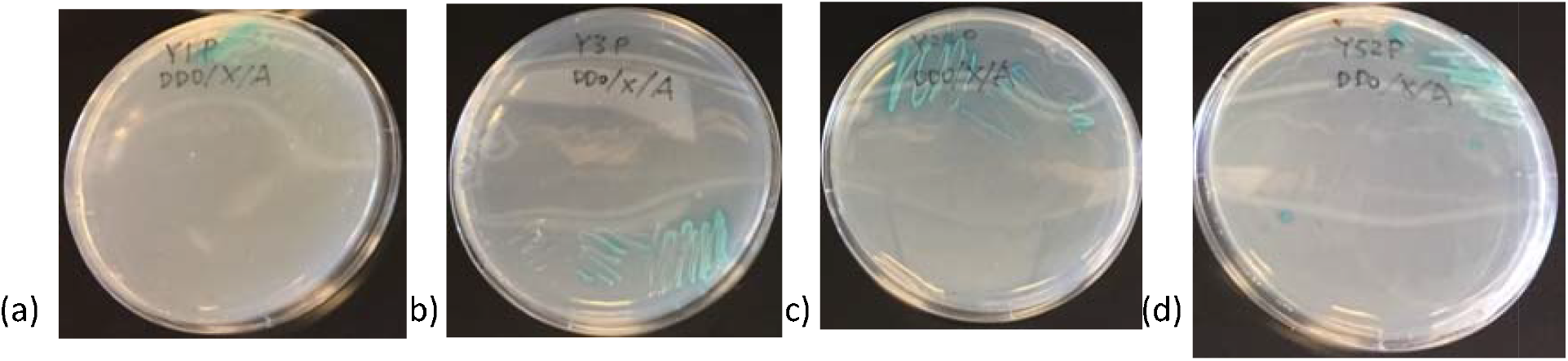
Viable diploid yeast colonies growing on SD/-Trp/-Leu/-His/-Ade/X-α-Gal/AbA (QDO/X/A) media (A) Positive clone #1 (B) Positive clone #3 (C) Positive clone #24 (D) Positive clone #52

### Co-immunoprecipitation to confirm novel protein interactions with PGRP3

To verify the results of our yeast two-hybrid screen, we selected two identified proteins i.e., MBNL3 and RBP5, to perform co-expression co-immunoprecipitation with PGRP3. Forward IP was done with Anti-His agarose to pull down PGRP3 protein complexes. Western blot was done on pull down elute to test for MBNL3 or RBP5 (Fig 3). Protein bands associated with MBNL3 and RBP5 were both present in the pull down fraction, confirming their interactions with PGRP3. A reverse co-immunoprecipitation was also completed with anti-Myc agarose. Subsequent western blot detected a protein band associated with PGRP3, thus corroborating our findings that MBNL3 and RBP5 interact with PGRP3 (Fig 4).

**Figure 3.**
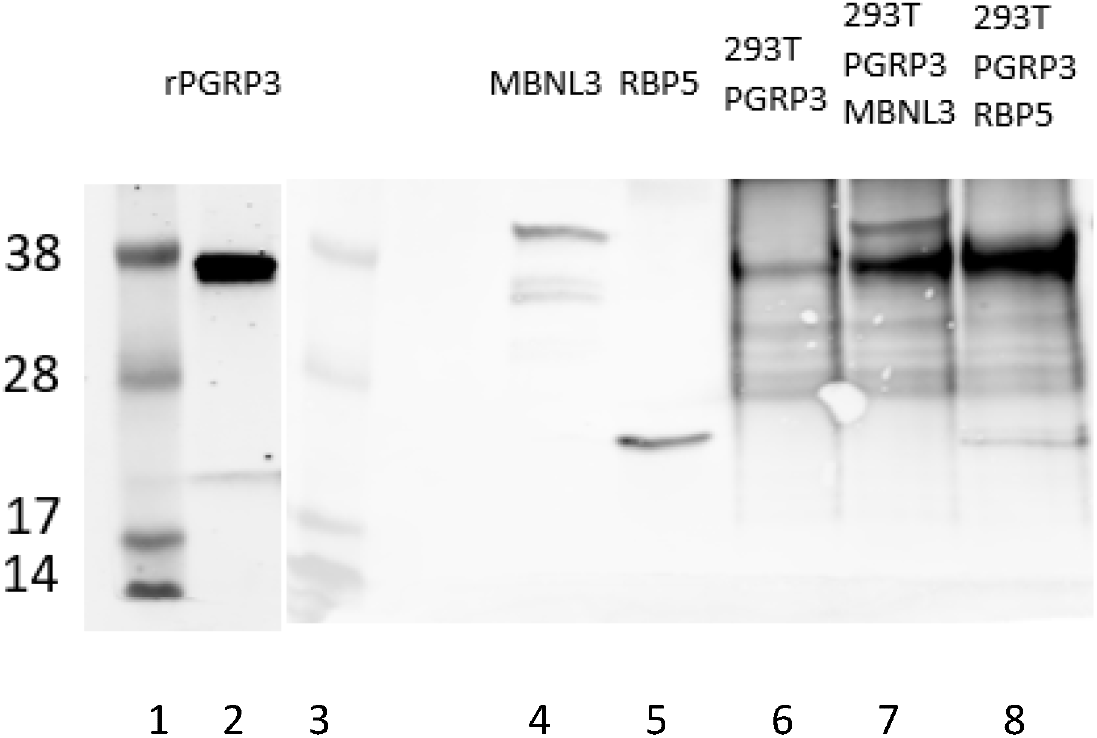
Western blot of PGRP3, MBNL3, and RBP5 using Myc staining. Lane 1 – ladder, Lane 2 – PGRP3 expression control, Lane 3 – ladder, Lane 4 – MBNL3 expression control, Lane 5 RBP5 expression control, Lane 6 −PGRP3 co-IP control, Lane 7 – MBNL3 co-IP, Lane 8 – RBP5 co-IP

**Figure 4.**
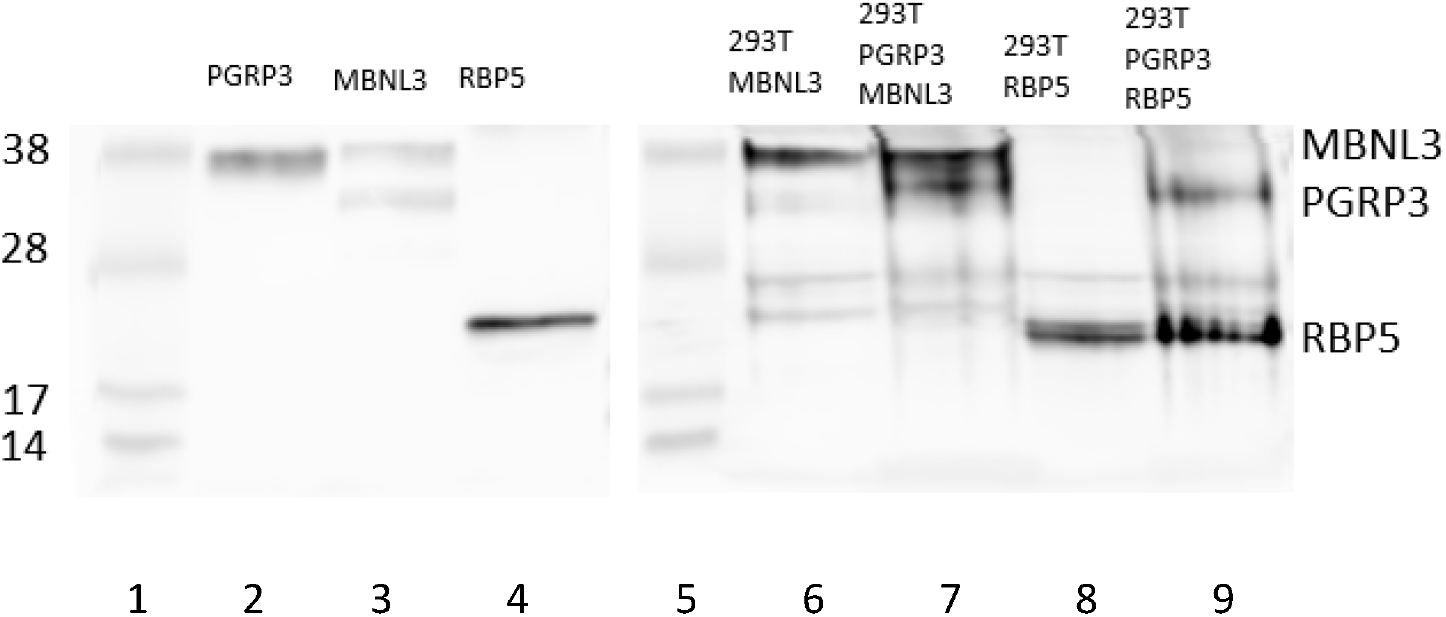
Western blot of PGRP3, MBNL3, and RBP5 using His staining. Lane 1 – ladder, Lane 2 – PGRP3 expression control, Lane 3 – MBNL3 expression control, Lane 4 – RBP5 expression control, Lane 5 – ladder, Lane 6 – MBNL3 co-IP control, Lane 7 – PGRP3 co-IP, Lane 8 RBP5 co-IP control, Lane 9 PGRP3 co-IP

## Discussion

Peptidoglycan recognition proteins (PGRP) are emerging as crucial mediators of tissue homeostasis. These secreted pattern recognition receptors surveil the environment and discriminate normobionts from pathobionts. PGRPs were first identified in the hemolymph of the *Bombyx mori* silkworm. They were described as proteins that could bind bacterial peptidoglycan and activate the prophenoloxidase cascade, which is involved in wound healing and melanization.^11^ In other insects like *Drosophila melanogaster*, PGRP activates the Toll and immune deficiency signal transduction pathways to create antimicrobial products, hydrolyze peptidoglycan, and induce phagocytosis.^12^ Further studies later revealed mouse and human PGRP orthologs, suggesting that the protein is highly conserved between insects and mammals.^13^

PGRP3 contains two PGRP domains that each have a single binding site for peptidoglycan.^14^ The crystal structure of PGRP3 has been reported to include three α helices and a central β-sheet consisting of five β-strands (four parallel and one anti-parallel). Its tertiary structure creates a fold that presumably provides two sites for molecular interaction—one that binds peptidoglycan and one that accommodates host effector or signaling molecules.^15,16^ The peptidoglycan binding groove is generally hydrophilic and filled with water molecules that are displaced following peptidoglycan binding.^16^ After peptidoglycan binds, a conformational change occurs in the PGRP domain that retains it in the binding groove.^17^ PGRP3 may form homodimers or heterodimers with PGRP4, which expands its specificity to peptidoglycan. Opposite the peptidoglycan binding sites are hydrophobic grooves that are thought to serve as binding sites for signaling molecules.^15^

In this yeast two-hybrid screen, we identified 4 potential proteins that interact with PGRP3: MBNL3, RBP5, ATXN2, and GPATCH8. Positive clones were first selected under SD/-Trp/-Leu/X-α-Gal/AbA (DDO/X/A) and then selected under SD/-Trp/-Leu/-Ade/-His/X-α-Gal/AbA (QDO/X/A) for higher stringency. Co-immunoprecipitation was able to confirm the interactions of MBNL3 and RBP5 with PGRP3. Extrapolating data from other studies regarding these 4 proteins may elucidate the biological pathways that PGRP3 is involved with in periodontitis.

MBNL3 is a one of three proteins in the human Muscleblind family of tissue-specific alternative splicing regulators. Alternative splicing is a process by which pre-mRNA transcripts are modified to include or exclude specific exons or noncoding regions. Different splicing patterns of the same gene may result in multiple protein isoforms with different functions. MBNL3 is most notably associated with the development of myotonic dystrophy, a disease characterized by gradual worsening muscle loss and weakness. It acts by inhibiting the expression of muscle differentiation markers like myogenin and myosin heavy chain.^18^ Primary splicing targets of MBNL3 include cardiac troponin T (cTNT) and insulin receptor (IR).^19^ A genome-wide study on peroxisome proliferator response elements (PPRE) identified these motifs in MBNL3, suggesting its regulation by PPARγ.^20^ Additionally, repression of MBNL3 has been correlated with neuraminidase-induced immune responses related to dendritic cells.^21^

RBP5 is a member of the lipocalin superfamily that binds vitamin A. Because mammals cannot make vitamin A, it must be obtained from food. Having RBPs bind to fat-soluble vitamin A increases its plasma concentrations 1,000-fold higher and also protects it from enzymatic and oxidative damage.^22^ Vitamin A plays a role in vision, embryonic development, immunity, and maintenance of epithelial surfaces.^23^ Besides acting as a carrier protein for vitamin A, RBP5 appears to have a role in preventing the progression of hepatocellular carcinoma. RBP5 levels were shown to be significantly downregulated in hepatocellular carcinoma tissue when compared to healthy liver tissues.^24^ There is also evidence that RBP5 has a role in lipid metabolism and is regulated by PPARγ in adipose tissue. Treatment of adipocytes with PPARγ agonist rosiglitazone significantly upregulated RBP5 expression via functional PPRE present in its promoter.^25^

ATXN2 is a polyglutamine protein whose dysfunction is commonly associated with spinocerebellar ataxia type 2, a disorder characterized by progressive loss of movement and coordination. The protein contains Lsm and LsmAD domains, which are highly conserved in RNA metabolism proteins.^26^ With these domains it may bind several RNA-binding proteins including RNA-binding motif protein 9 (RBM9) and RNA-binding protein with multiple slicing (RBPMS).^27^ ATXN2 may also upregulate translation through polyribosome and Poly(A)-binding protein (PABP) or downregulate translation through the formation of stress granules.^28^ Studies have also demonstrated ATXN2’s participation in biological processes such as actin cytoskeleton reorganization and endocytic vesicle assembly.^29,30^ Through interaction of its proline-rich domains with SH3 motifs in tyrosine protein kinase Src and Growth factor receptor-bound protein 2 (Grb2), ATXN2 has been shown to inhibit endocytosis machinery.^31^

GPATCH8 is a protein whose function has not been explored as extensively. Missense mutations in the GPATCH8 gene have been linked to hyperuricemia cosegregating with osteogenesis imperfecta.^32^ In a different study, computations by NCBI’s Conserved Domain Architecture Retrieval Tool (CDART) predicted that GPATCH8 contains a KOG0106 domain, which is part of the SRp55/B52/SRp75 superfamily and known to facilitate alternative splicing. This study also used the Simple Modular Architecture Research Tool (SMART) to identify a Cys_2_His_2_ (C2H2) zinc finger domain in GPATCH8, further substantiating evidence for its RNA splicing activity.^33^

PGRP3 binding to these proteins posits a significant role in modulating the cellular functions of these protein partners. Hence, we speculate that PGRP3 may play a role in RNA splicing and processing due its ability to bind MBNL3, ATXN2, and GPATCH8. An additional theme that is common to PGRP3 binding proteins is endocytic trafficking as is known for ATXN2 and RBP5. An intriguing observation is that MBNL3, RBP5, and the already identified DHTKD1 each contain similar PPRE motifs in their gene promoters as PGRP3. PGRP3 expression has previously been shown to be induced by PPARγ ligands, leading to downregulation of inflammatory cytokines IL-8, IL-12, and TNF-α. Functional protein studies will need to be done to understand how PGRP3 interactions alter these cellular processes.

Through this study we have identified some novel PGRP3 interactions. Given the structure of PGRP3 there are likely other interactions that need to be identified. Although one of the strengths of a yeast two-hybrid screen is its ability to filter through millions of proteins, one of its main drawbacks is the number of false positives and false negatives that are produced. False positives are less of an issue because they can be confirmed as genuine protein interactions with other assays. False negatives, however, present a problem in developing a protein interactome because they limit our ability to detect every protein that binds to PGRP3. For instance, our yeast two-hybrid screen did not select for the previously reported PGRP-interacting proteins DHTKD1 and MICA. It conceivable that this is likely due to the requirement of Y2H screens that interactions must occur in the nucleus of yeast cells, resulting in failure to capture interactions that occur with the secreted PGRP3 in the extracellular matrix.

## Conclusions

In this study, we have identified novel interacting partners of PGRP3, namely ATXN2, MBNL3, GPATCH8, and RBP5. ATXN2, MBNL3, and GPATCH8 are known to be critical for RNA processing. RBP5 has known transporter activity, while ATXN2 is also known for cytoskeleton reorganization and endocytic trafficking. We also note the intriguing observation that a majority of PGRP3 interacting proteins are under the transcriptional control of PPARy, suggesting these interactions may be biologically meaningful. This premise needs to be further explored with functional studies.

**Table 2.** Genes identified from yeast two-hybrid screen and their corresponding gene ID, description, chromosome location, and protein size

| Symbol | Gene ID | Gene Description | Chromosome Location | Protein Size |
| --- | --- | --- | --- | --- |
| MBNL3 | 55796 | Muscleblind-like Protein 3 | Xq26.2 | 38 kDa |
| RBP5 | 83758 | Retinol Binding Protein 5 | 12p13.31 | 15.9 kDa |
| ATXN2 | 6311 | Ataxin 2 | 12q24.12 | 140 kDa |
| GPATCH8 | 23131 | G Patch Domain-containing Protein 8 | 17q21.31 | 164 kDa |

